# Competitive interaction between ATP and GTP regulates mitochondrial ATP-sensitive potassium channels

**DOI:** 10.1101/2023.01.24.525224

**Authors:** Plinio Bezerra Palácio, Geovanna Carvalho de Freitas Soares, Gabriella Moreira Bezerra Lima, Pedro Lourenzo Oliveira Cunha, Anna Lidia Nunes Varela, Heberty Tarso Facundo

## Abstract

Mitochondrial ATP-sensitive K^+^ channels (mitoKATP) have been recently characterized structurally, and possess a protein through which K^+^ enters mitochondria (MitoKIR), and a regulatory subunit (mitoSUR). The mitoSUR regulatory subunit is an ATP-binding cassette (ABC) protein isoform 8 (ABCB8). Opening of these channels is known to be cardioprotective, but the molecular and physiological mechanisms that activate them are not fully known. Here, to better understand the molecular and physiological mechanisms of activators (GTP) and inhibitors (ATP) on the activity of mitoKATP, we exposed isolated mitochondria to both nucleotides. We also used molecular docking directed to the nucleotide-binding domain of human ABCB8/mitoSUR to test a comparative model of ATP and GTP effects. As expected, we find that ATP dose-dependently inhibits mitoKATP activity (IC_50_ = 21.24 ± 1.4 mM). However, simultaneous exposure of mitochondria to GTP dose-dependently (EC_50_ = 13.19 ± 1.33 mM) reversed ATP inhibition. Pharmacological and computational studies suggest that GTP reverses ATP activity competitively. Docking directed to the site of crystallized ADP reveals that both nucleotides bind to mitoSUR with high affinity, with their phosphates directed to the Mg^2+^ ion and the walker A motif of the protein (SGGGKTT). These effects, when combined, result in GTP binding, ATP displacement, mitochondrial ATP-sensitive K^+^ transport, and lower formation of reactive oxygen species. Overall, our findings demonstrate the basis for ATP and GTP binding in mitoSUR using a combination of biochemical, pharmacological, and computational experiments. Future studies may reveal the extent to which the balance between ATP and GTP actions contribute toward cardioprotection against ischemic events.

## 1. Introduction

The mitochondrial ATP-sensitive K^+^ channel (mitoKATP) drives K^+^ into the mitochondrial matrix controlling organelle volume [1,2]. This channel is negatively modulated by adenine nucleotides (ATP and ADP) and positively by guanine nucleotides (GTP and GDP) [3,4]. Little is known about the binding and action of nucleotides in the mitoKATP regulatory protein. MitoKATP controls mitochondrial volume more significantly than membrane potential [1], as well as having effects modulating Ca^2+^ overload [5,6] and reactive oxygen species (ROS) formation [7–9]. Similar to KATP channels at the plasma membrane, the mitochondrial channel is a target for the antidiabetic sulfonylurea drug glibenclamide (that inhibits K^+^ currents) and diazoxide (a mitoKATP opener) [7,10–14]. It is important to point out that even though K^+^ currents through a mitoKATP have been described since the early 1990s [15] its molecular composition was unknown until recently, when an elegant study demonstrated that mitoKATP is composed of a pore-forming protein (CCDC51) and a regulatory subunit (ABCB8 – mitoSUR) [2]Nevertheless, the molecular structure of the pore-forming channel is still not resolved. Interestingly, the regulatory subunit (mitoSUR) has been independently crystallized in the presence of ADP (available at https://www.rcsb.org/under code 5OCH) or the presence of the ATP analog adenylyl-imidodiphosphate (AMPPNP-available at https://www.rcsb.org/ under code 7EHL). A typical ABC transporter protein, ABCB8 is composed of a transmembrane domain (TMD) and a nucleotide-binding domain (NBD) [16].

MitoKATP opening has been linked to ischemia/reperfusion protection triggered by ischemic preconditioning (induced by short cycles of ischemia) or pharmacological preconditioning [7,10,17,18]. Papers usually explore the pharmacology of mitoKATP openers and blockers but neglect the impact of physiological regulators. As this channel is modulated negatively by adenine (ATP and ADP) and positively by guanine nucleotides (GDP and GTP) [3,4,8], it is plausible that these nucleotides are associated with endogenous activation or blockage of this channel (under physiological and pathological conditions). The positive effects of GTP on mitoKATP opening are blocked pharmacologically by glibenclamide and 5-hydroxydecanoate [19]. Most importantly, mitoKATP activity blocks cell death, excessive reactive oxygen species (ROS) formation, and Ca^2+^ dysregulation under pathological conditions [7–9]. The regulatory subunit of mitoKATP (mitoSUR/ABCB8) is implicated in mitochondrial iron export, and its disruption leads to cardiomyopathy [20]. Previously, we found that glibenclamide or 5-hydroxidecanoate, two inhibitors of mitoKATP, prevented the beneficial effects of mitoKATP opening, leading to cardiac hypertrophy [11,13].

The development of more specific mitoKATP openers or blockers requires a better understanding of how the channel works physiologically, how endogenous compounds modulate it, and the resulting consequences. In the present study, we investigated the effects of nucleotides (GTP and ATP) on mitoKATP in isolated mitochondria. Our results clearly show that GTP reverses ATP-inhibited mitochondrial K^+^ transporte. To understand the underlying mechanisms, we computationally docked both nucleotides to the NBD (directed to the same site as crystallized ADP) of mitoSUR (ABCB8). From these in silico simulations and additional experimental data, we propose that GTP opens mitoKATP by modulating ATP binding, which is suggestive of competitive antagonism. In addition, our studies have shown and confirmed that GTP can prevent ROS production induced by mitoKATP inhibition, contributing to a debated point in the literature regarding mitoKATP.

## 2. Materials and Methods

All aqueous solutions were prepared in deionized water. Rotenone, and oligomycin were dissolved in dimethyl sulfoxide (DMSO). GTP, ATP and succinate solutions were prepared in water and corrected to a pH value between 7.0 and 7.4 with NaOH. All chemicals were purchased from Sigma-aldrich.

### 2.1. Animals

All procedures were approved by the Institutional Animal Experimentation Ethics Committee of the Universidade Federal do Cariri (protocol number 02/2020). All experiments were performed according to the Guide for the Care and Use of Laboratory Animals published by the National Institutes of Health. 60-day-old Swiss male mice weighing 25–30 g were anesthetized with ketamine (100 mg/kg) and xylazine (10 mg/kg) before experiments.

### 2.2. Isolation of cardiac mitochondria

Mitochondria were isolated from male mouse cardiac tissue. Mice were anesthetized, and their hearts were immediately removed and washed in an ice-cold buffer containing 300 mM sucrose, 10 mM K^+^ HEPES buffer, and 1 mM K^+^ EGTA buffer (isolation buffer) at pH 7.2. The tissue was finely minced and incubated in protease type I (Sigma-Aldrich) for 10 minutes. Then, the protease was washed away with the same solution, with 1 mg/mL BSA added. Cardiac tissue was homogenized using a glass homogenizer (potter). Then, nuclei and cellular residues were pelleted by centrifugation (5 minutes) at 600 g. The last supernatant was recentrifuged at 9400 g for 8 minutes to obtain a mitochondrial pellet. Finally, the mitochondrial pellet was resuspended in isolation buffer (100–150 L). Samples were kept on ice and used within 1 hour of isolation to ensure mitoKATP activity. All mitochondrial experiments were conducted using 50–100 mg of protein.

### 2.3. Mitochondrial ATP-sensitive, K^+^- dependent, swelling

Mitochondria (50 μg protein) were incubated in an experimental buffer containing: 100 mM KCl, 10 mM HEPES, 4 mM succinate, 1 μM rotenone, 2 mM MgCl2, 2 mM KH_2_PO_4_, and 1 μg/mL oligomycin, pH 7.2 (KOH). We collected light-scattering traces within 1 hour of mitochondrial isolation to ensure mitoKATP activity. Mitochondrial ATP-sensitive K+ entrance and swelling were evaluated over time using a spectrophotometer at 540 nm. The delta of absorbance was calculated as the difference between 0 and 90 seconds after the beginning of each trace.

### 2.4. Mitochondrial respiration

Mitochondrial O_2_ consumption was determined using a Clark-type electrode (Hansatech, UK), at room temperature. The mitochondrial suspension (100 μg protein) was incubated in a reaction buffer consisting of 100 mM KCl, 10 mM HEPES, 2 mM MgCl_2_, 2 mM KH_2_PO_4_, and 4 mM succinate plus rotenone (1 μM) pH 7.2 (KOH). State 3 (ADP-dependent oxygen consumption) was initiated with 1 mM ADP. State 4 was induced with oligomycin (1 μg/mL). Respiratory control ratios (RCR) were calculated as state 3/state 4. Uncoupled respiration was induced with CCCP (100 nM).

### 2.5. H_2_O_2_ Measurements

*Mi*tochondrial H_2_O_2_ production was detected by incubating cardiac mitochondria (protected from light) with Amplex red (50 μmol/L) and horseradish peroxidase (1 U/mL) at 37°C for 60 min at pH 7.2. The reaction buffer consisted of 100 mM KCl, 10 mM HEPES, 2 mM MgCl_2_, 2 mM KH_2_PO_4_, 1 μg/mL oligomycin, and 4 mM succinate plus rotenone (1 μM) pH 7.2 (KOH). After 1 hour, the tubes were centrifuged at 9300 g for 2 minutes. The supernatant was transferred to a cuvette, and the absorbance was measured at 560 nm. Amplex red and horseradish peroxidase were incubated without the sample to determine background absorbance. H_2_O_2_ released was calculated in μmol/mg protein using a calibration curve created using H_2_O_2_ standards.

### 2.6. Molecular docking

The X-ray crystal structure of human ABCB8 protein (PDB code 5OCH) was downloaded from the protein data bank website (PDB) (https://www.rcsb.org/structure/5OCH) and used for molecular docking using the autodock Vina built-in Chimera modeling software from the University of California, San Francisco (UCSF). The chemical structures of compounds were imported into the Chimera software by the canonical smiles found on https://pubchem.ncbi.nlm.nih.gov and used as mol2 files for docking. All crystallographic water and small molecules were removed. We computationally reversed YCM483 back to cysteine using UCSF Chimera. The macromolecules were prepared with Gasteiger partial charges, and hydrogens were added. The nucleotides (ATP and GTP) docking to the protein (ABCB8) focused in on the nucleotide-binding site of ADP (x = 203.89, y = 8.86, and z = 436.92). The 2D and 3D interactions were imported into the Discovery Studio visualizer, which was used to identify significant interactions between the ligands (GTP and ATP) and the receptor protein. The dissociation constant (Kd) was calculated from experimental binding free energies using in-house developed Python-based software, applying the equation:

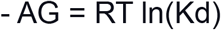

where R is the gas constant (1.98 cal/K mol) and T is the temperature (298 K).

#### Pharmacological test for competitive inhibition

The docking results prompted us to attempt to pharmacologically characterize the interaction between GTP or ATP with mitoKSUR (ABCB8). We used a concentration-response curve of GTP in the presence of increasing doses of ATP (100, 200, and 500 mM). Then we determined the EC_50_ of the GTP in each curve. Importantly, we could not run the test in ATP`s absence since in these settings the channel is already open. GTP causes dose-dependent mitochondrial swelling secondary to mitochondrial K^+^ entrance.

### 2.12. Statistical analysis

We used Graphpad Prism software for statistical analysis. The data are presented as mean ± S.E.M. Analysis was conducted using ANOVA followed by a Tukey’s test. P < 0.05 was regarded as statistically significant. All vector graphics and dose-response curves were created using GraphPad Prism software.

## 3. Results

### 3.1. ATP binding inhibits mitochondrial ATP-sensitive K^+^ entrance

ATP inhibits mitochondrial ATP-sensitive K+ entrance at micromolar concentrations. MitoKATP is also sensitive to guanosine nucleotides (GDP and GTP) [14]. Some studies point out that ATP binds to a mitochondrial sulfonylurea receptor, recently characterized as the ABCB8 protein (mitoSUR) [2]. This channel is also inhibited by ADP and activated by either GDP or GTP [3,4,8]. The importance of mitoKATP is underscored by the observation that pharmacological inhibition of mitoKATP worsens the outcome in cardiac tissue undergoing ischemia and reperfusion [7,10,12] and cardiac hypertrophy [11,13]. However, how mitoKATP binds and senses changes in ATP or other nucleotide (GTP, GDP) concentrations is still unclear. Particularly, mitochondria produce approximately 95% of the ATP in cardiomyocytes, cells in which ATP levels are usually high [21]. This simple observation would challenge the importance of mitoKATP opening under physiological conditions. Therefore, we aimed to understand the nucleotide (ATP and GTP) regulation of mitoKATP. We began by testing the effects of ATP on mitoKATP. First, we exposed isolated cardiac mitochondria to different concentrations of ATP and tested them for swelling secondary to K^+^ uptake. In the absence of ATP, isolated cardiac mitochondria reduced light scattering (indicating swelling and mitoKATP activity). As expected, ATP blocked it dose-dependently (Fig. 1). We calculated the ATP binding IC_50_ in the micromolar range of 21.24 ± 1.4 μM.

**Figure 1.**
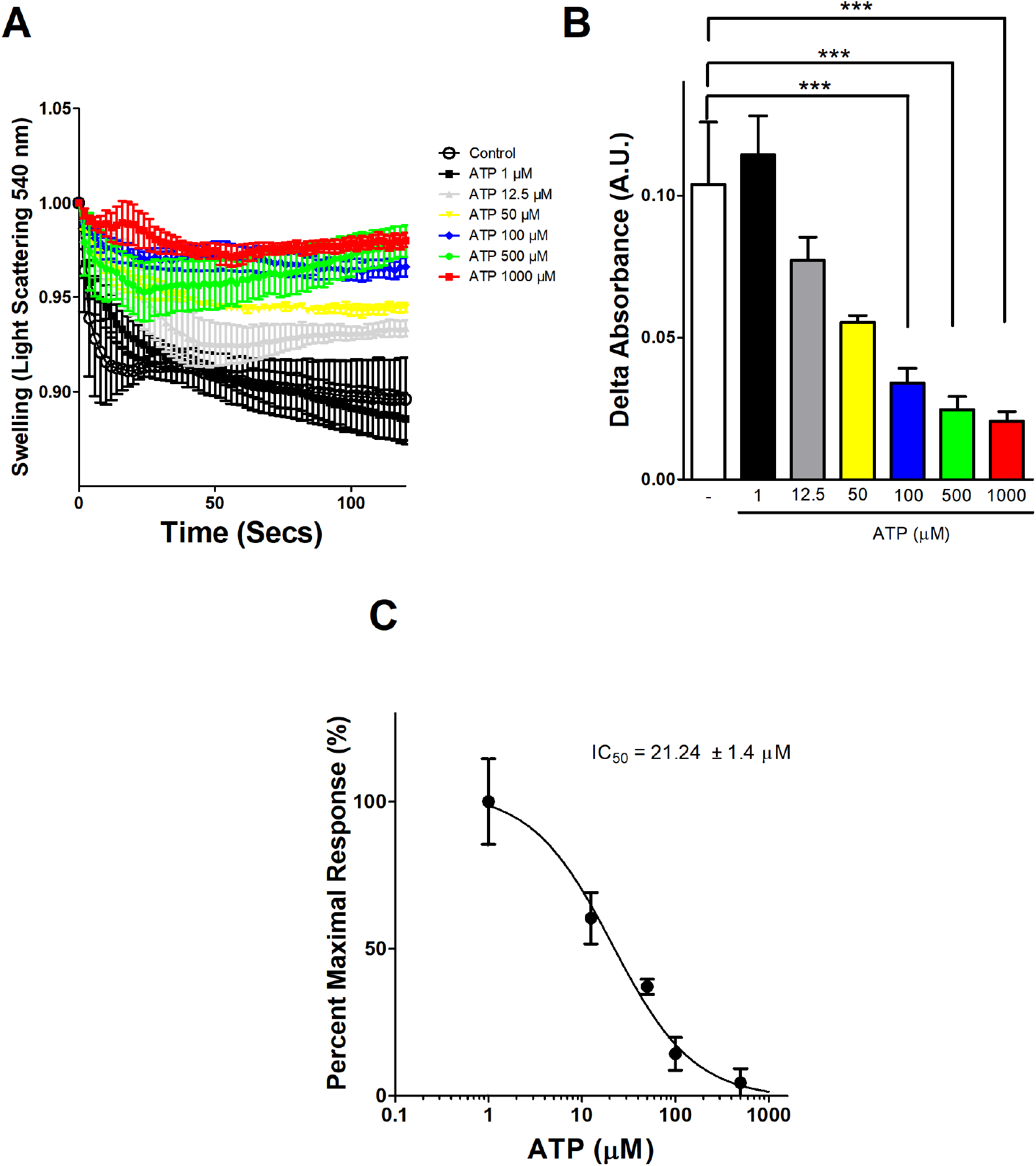
Mitochondrial K^+^ entrance is dose-dependently inhibited by ATP. (A) Light scattering traces of mitochondria (50 μg protein/mL) incubated in 100 mM KCl, 10 mM HEPES, 2 mM succinate, 1 μM rotenone, 2 mM MgCl_2_, 2 mM KH_2_PO_4_, 1 μg/mL oligomycin, pH 7.2 (KOH). Representative traces of mitochondrial swelling with no further additions (control) and in the presence of increasing ATP concentrations 1-1000 μM. (B) represents averages ± S.E.M of changes in light scattering calculated after 90 seconds of incubation, as in **(A)**. (C) Dose-response of ATP sensitive mitochondria K^+^ entrance. * P< 0.05, *** P<0.001, n = 4 per group.

### 3.2. GTP binding activates mitochondrial ATP-sensitive K^+^ entrance

As stated before, mitoKATP is susceptible to nucleotides such as ATP, ADP, GTP, and GDP. GTP activates mitoKATP in a manner sensitive to pharmacological inhibitors, such as 5-hydroxydecanoate and glibenclamide [4,14]. Here, we conducted additional experiments in isolated mitochondria to test the positive effects of GTP on mitoKATP. These experiments revealed that increasing GTP (or the absence of ATP - control trace) opens mitoKATP, leading to mitochondrial K^+^ and H2O uptake (Figure 2A,B). These two events combined will lead to a new mitochondrial volume steady-state. Indeed, GTP opens mitoKATP in a dose-dependent manner (EC_50_ = 13.19 ± 1.33 mM), even at high ATP (500 mM, Figure 2B,C). In summary, GTP opens mitoKATP dose-dependently (Fig. 2), blocking ATP inhibitory effects.

**Figure 2.**
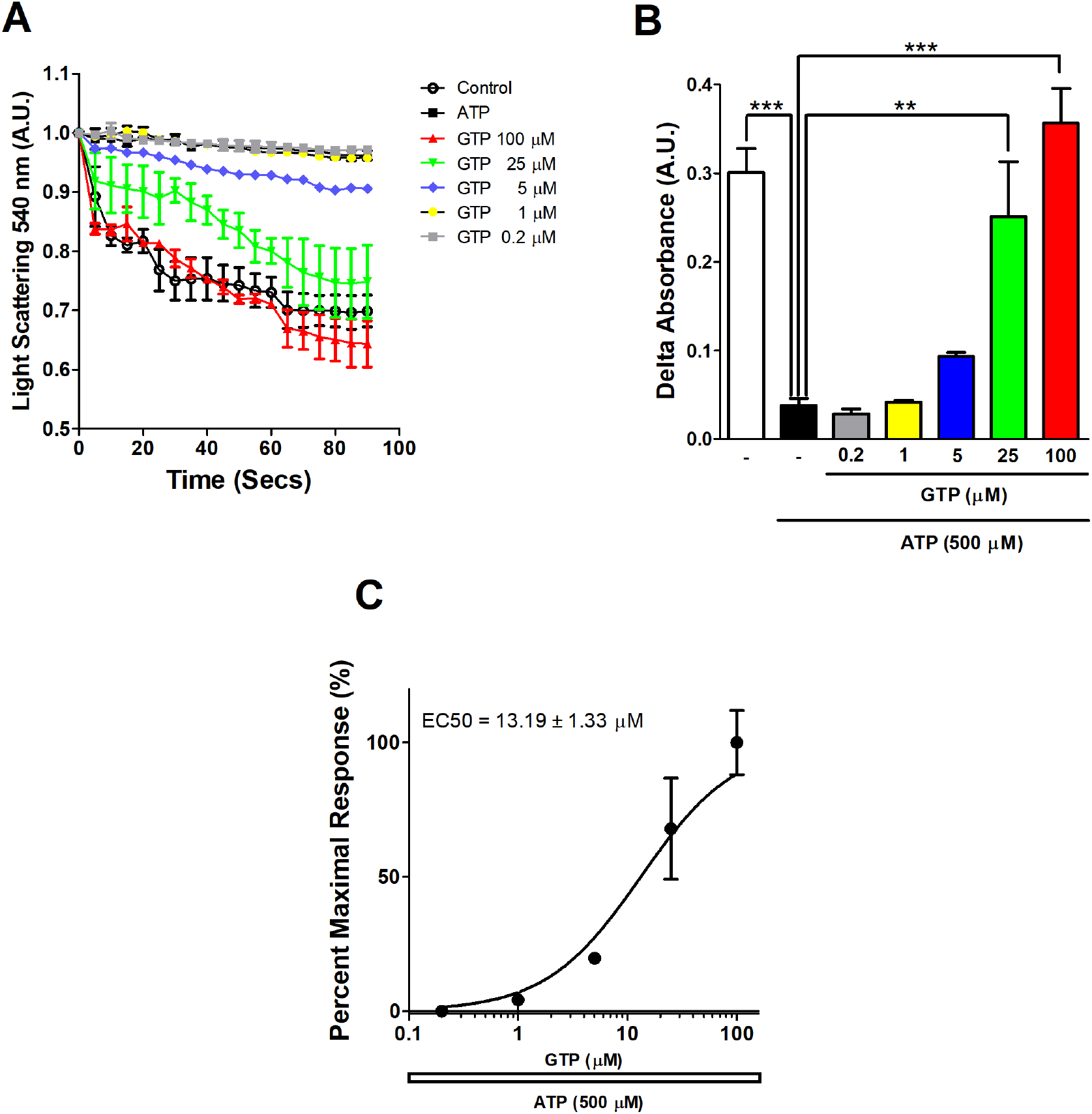
GTP dose-dependently activates mitochondrial ATP-sensitive K^+^ entrance. (A) Light scattering traces of mitochondria (50 μg protein/mL) incubated in 100 mM KCl, 10 mM HEPES, 2 mM succinate, 1 μM rotenone, 2 mM MgCl_2_, 2 mM KH_2_PO_4_, 1 μg/mL oligomycin, pH 7.2 (KOH), plus 500 μM ATP. Average ± S.E.M traces of mitochondrial swelling with no further additions (control) and in the presence of increasing GTP concentrations 0.2 -100 μM. (B) represents averages ± S.E.M of changes in light scattering calculated after 90 seconds of incubation, as in (A). (C) Dose-response of GTP effects on mitochondrial ATP-sensitive K^+^ entrance. * P< 0.05, *** P<0.001, n = 4 per group.

### 3.3. Competitive ABCB8 (mitoSUR) binding between GTP and ATP

We decided to study the binding of ATP to the ABCB8 protein (also known as mitoSUR) using molecular docking. To accomplish this, we downloaded the X-ray crystal structure of human ABCB8 from the protein data bank (PDB code 5OCH) and used it for docking at the same site as crystallized ADP in the structure (Fig. 3A, as described in materials and methods). The best interaction between ABCB8 and ATP gave a negative free binding energy of -8.5 kcal/mol. ATP binds firmly at the target site of mitoSUR with fourteen conventional hydrogen bonds (Table I). Additionally, this interaction resulted in a calculated dissociation constant of 0.554 mM (Table I). The ATP beta and gamma phosphates are directed toward the Mg^2+^ ion inside the Walker A motif (Fig. 3A). The Mg^2+^ ion electrostatically interacts with oxygen in the ATP beta and gamma phosphates. The oxygens in gamma phosphate also interact electrostatically and form hydrogen bonds with LYS513. A closer look at the interaction between ATP and ABCB8 revealed that the ATP alpha, beta, and gamma phosphates had nine predicted hydrogen bonds with residues TYR481, GLY510, GLY511, GLY512, and THR515 inside chain A of the protein. Many amino acids interact via Van der Walls (SER509, THR514, VAL489, GLY606, ARG608, THR610), carbon-hydrogen bonds (GLY512), conventional hydrogen-bonds (PHE487, GLU607, GLY609, THR611), pi-pi T-shaped (TYR481), and pi-alkyl (CYS483 and ARG484 - Fig. 3B).

**Figure 3.**
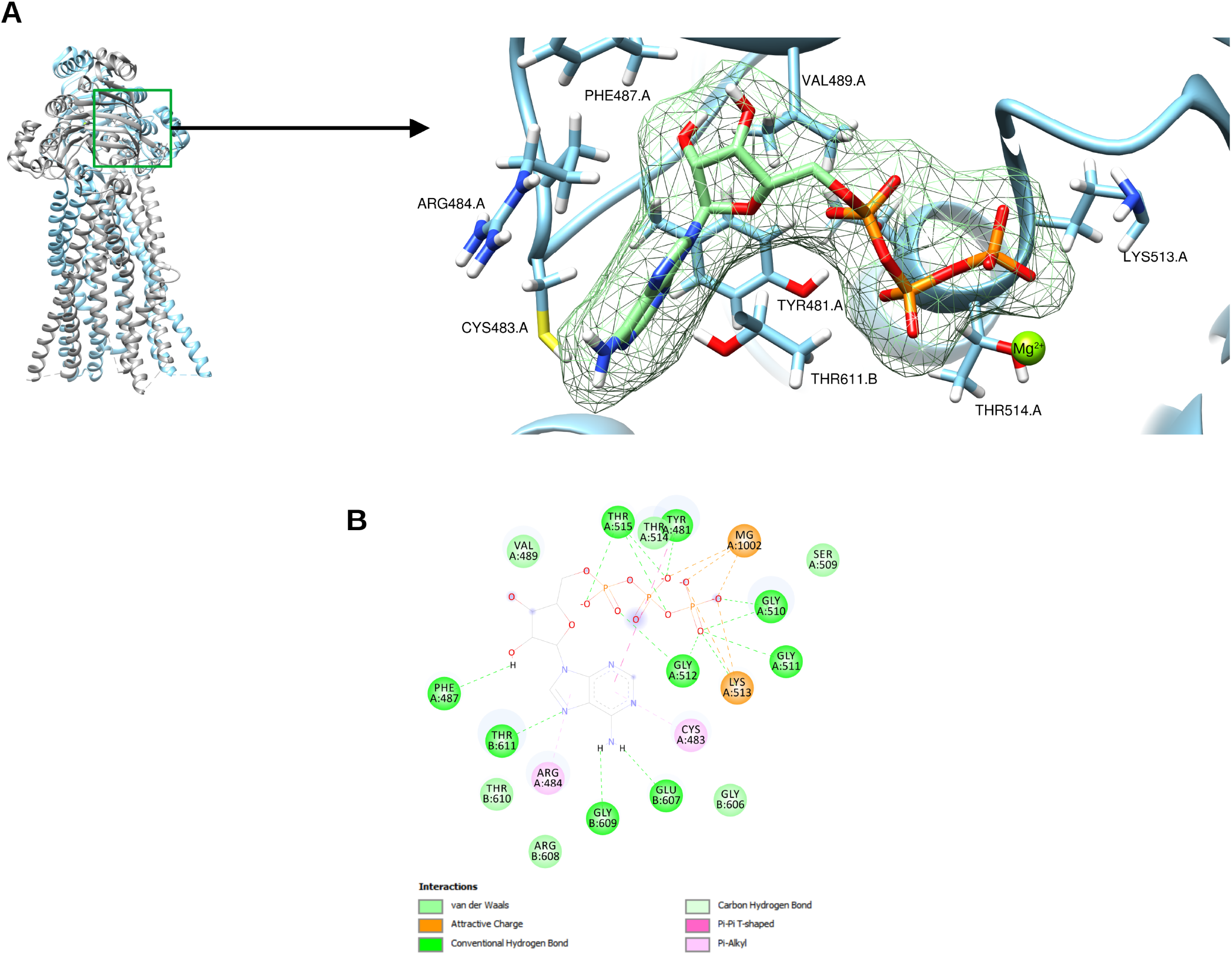
Docked pose of ATP on ABCB8 (mitoSUR). (A) Best ATP binding position in ABCB8 residues (B) A 2D representation showing the interactions of ATP and ABCB8 residues Hydrogen bonds are shown in green dashed lines. Orange dashed lines indicate critical attractive charges.

**Table 1:** Score for the best interaction pose seen in Fig. 5, calculated Kd and number of hydrogen bonds.

|  | **Binding Energy**<br>**Score (Kcal/mol)** | **Hbonds** | **Calculated Kd** |
| --- | --- | --- | --- |
| **ATP** | - 8.5 | 14 | 0.554 μM |
| **GTP** | - 8.1 | 6 | 1.29 μM |

Knowing that GTP activates mitoKATP even in the presence of elevated levels of ATP (Fig. 2), we conducted additional *in silico* experiments to simulate the mechanisms of GTP binding into the mitoKATP protein complex. To accomplish this, we directed the docking to the same region in which ADP is bound in the crystal structure of ABCB8, as depicted in the PDB website (nucleotide-binding site). The interactions between GTP and the ABCB8 dimer were similar to those seen for ATP docking (Fig. 4). The oxygens in GTP’s gamma phosphate interact with Lys513 via attractive charges. Mg^2+^ exhibits the same attractions as Lys513, but with the addition of beta-phosphate oxygen. This ion also interacts with the charged oxygen in the beta phosphate. The phosphates (alpha, beta, and gamma) are additionally held in place by hydrogen bonds. More specifically, GLY510, GLY511, GLY512, THR514, and THR515 residues interact with the oxygens in gamma phosphate. THR515 also interacts with the oxygen in the alpha phosphate. Many mitoSUR (ABCB8) amino acids interact with the GTP molecule via Van der Walls interactions (ASP244, TYR624, TYR481, VAL489, ARG342, ASP607, THR610, and THR611). An important observation here is that the best interaction between mitoSUR and GTP gave a negative free binding energy of -8.1 kcal/mol, from a total of 6 hydrogen bonds. Interestingly, this interaction resulted in a calculated dissociation constant of 1.29 mM (Table I). In parallel, we also found by computational analysis using GTP binder (https://webs.iiitd.edu.in/raghava/gtpbinder/algo.html) that ABCB8 binds GTP mainly in residues GLY510, GLY511, GLY512, K513, THR514 (results not shown). Taken together, these results suggest that a strong interaction is formed between mitoSUR and GTP. It is important to note that our models suggest a potential competitive inhibition between GTP and ATP. Taking this into account, we present a docking model of the suggested competitive inhibition, showing that the best binding positions of GTP and ATP are on similar amino acids in the ABCB8 protein nucleotide binding site (Fig. 5). Knowing that, we constructed a topological structure highlighting ABCB8 amino acids that interact with ATP and GTP. This visual methodological approach reveals an overlapping interaction between ATP and GTP (Supplementary Fig. 1). One could argue that GTP interacts with ASP244 and ARG342, which lie outside the nucleotide-binding domain. Indeed, both residues are located inside intracellular loops (coupling helixes) 1 and 2 (ICL1 and 2, respectively). It is important to point out that ARG342 interacts with the crystallized ADP molecule (PDB code 5OCH). Indeed, several residues interact with both GTP and crystallized ADP (Supplementary Table I). Knowing all that, we constructed a series of dose-response curves using GTP in the presence of increasing concentrations of ATP (100, 200, and 500 mM). These dose-responses show a clear shift to the right from 100 up to 500 mM ATP, and point to a competitive antagonism, as predicted in our docking, pharmacological studies, and elsewhere [4]. Interestingly, all aminoacids interacting with ATP and GTP are conserved among different species (Results not shown).

**Figure 4.**
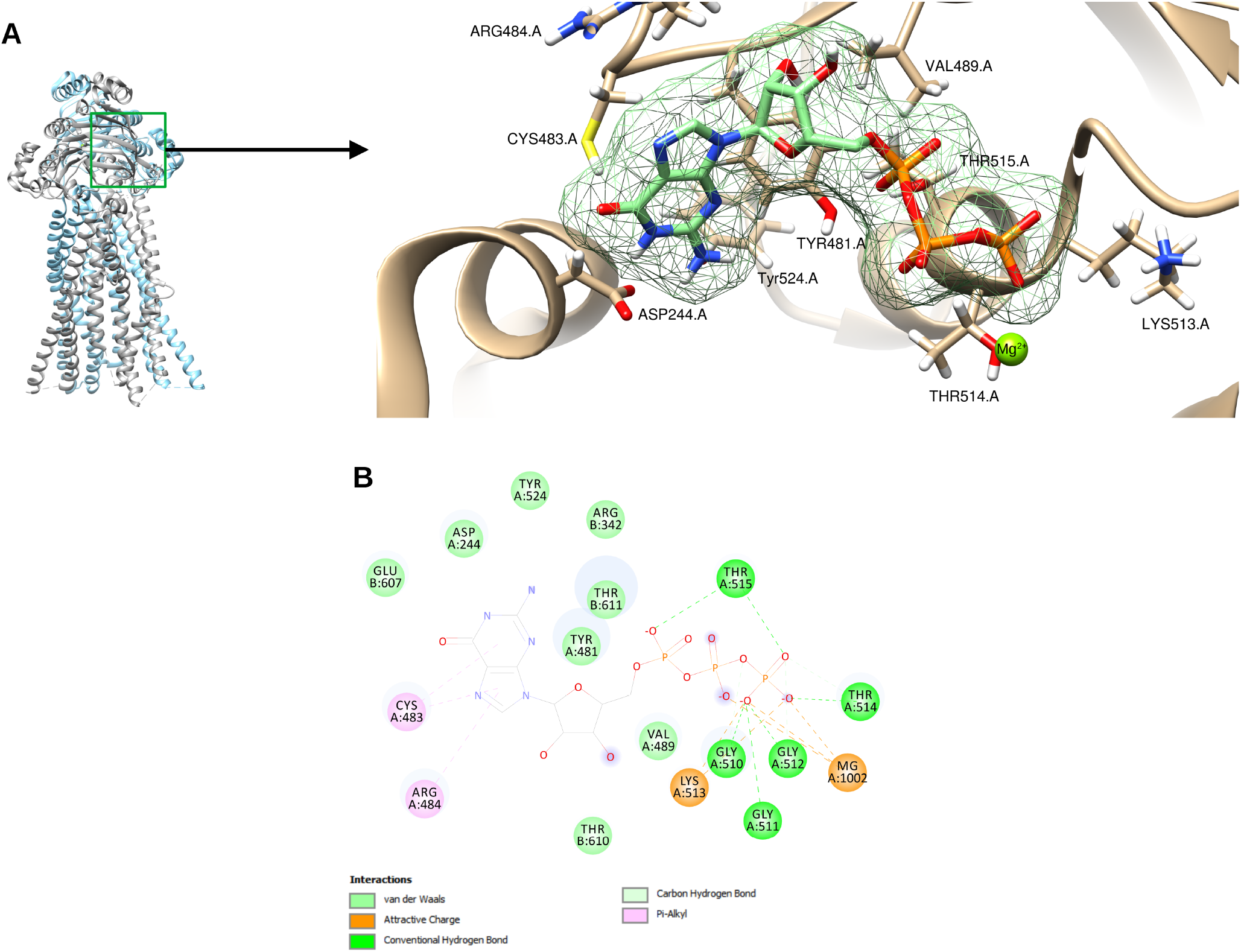
Docked pose of GTP on ABCB8 (mitoSUR). (A) Best GTP binding position in ABCB8/mitoSUR residues (B) A 2D representation of ATP and ABCB8/mitoSUR residue interactions Hydrogen bonds are shown in green dashed lines. Orange dashed lines indicate critical attractive charges.

**Figure 5.**
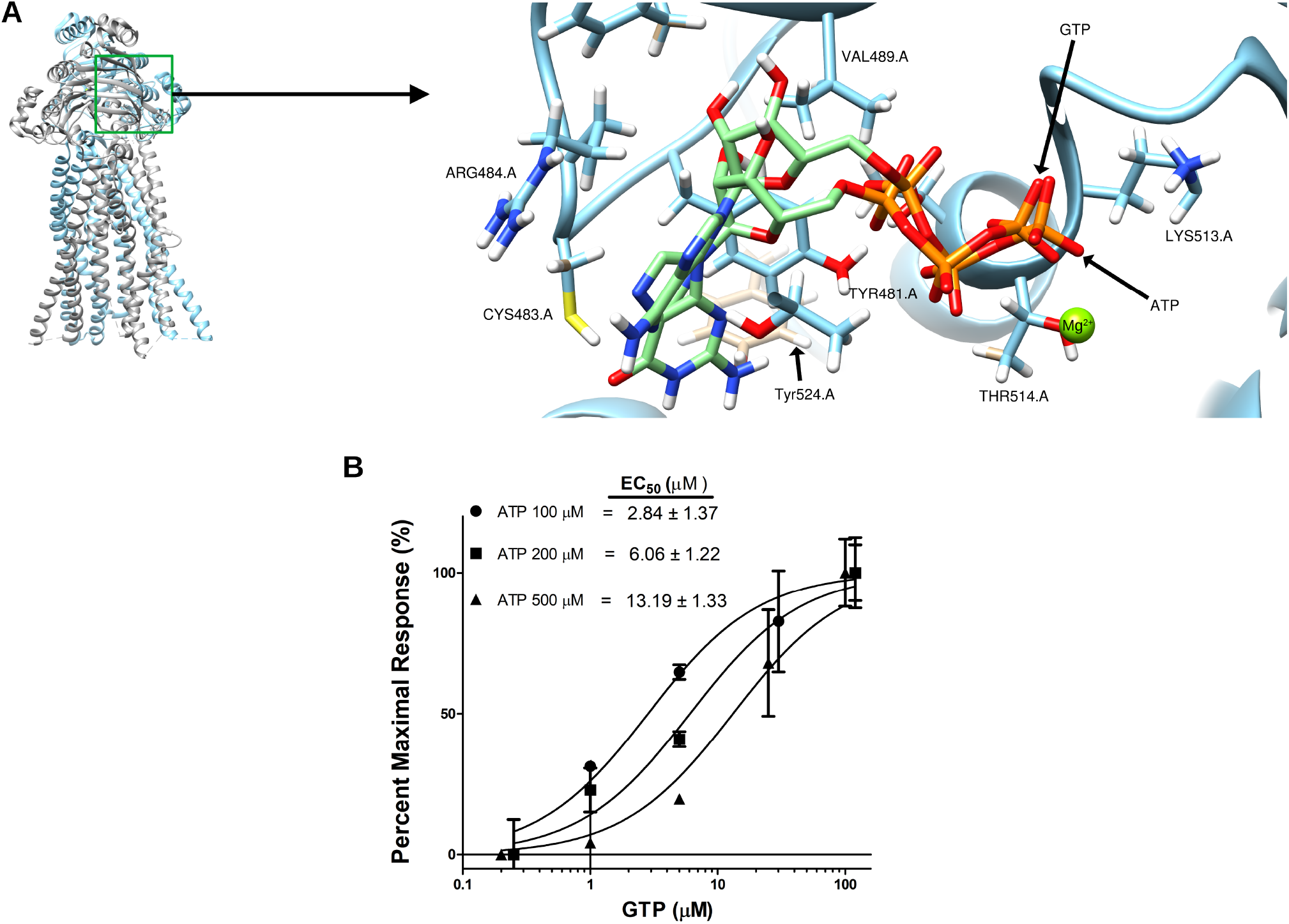
Pharmacological and nucleotide binding domain-directed docking reveal a possible competition between GTP and ATP for mitoSUR binding. A. Computational docking showing the best binding positions of GTP and ATP in the ABCB8/mitoSUR protein. The binding of ATP overlaps the GTP binding site. The diagrams were prepared using Chimera software. B. ATP-binding affects mitoKATP sensitivity to GTP. Dose-response curves of GTP (0.2 – 100 mM) in the presence of increasing concentrations of ATP (100, 200, and 500 mM). n = 4 per group.

### 3.5. ATP and GTP effects on mitochondrial ROS production

There is some controversy regarding if mitoKATP opening directly generates or blocks mitochondrial ROS formation, although it is clear that under pathological conditions, mitoKATP avoids the harmful overproduction of ROS [7,8,22]. Knowing that, we tested whether GTP-triggered mitoKATP opening modulates mitochondrial ROS production in the presence of low or high ATP levels. Mitochondria dynamically use substrates and consume oxygen, generating ATP. So, we also tested for the effects of GTP on mitochondrial O_2_ consumption. As seen in Fig. 6 (a representative trace), GTP does not cause any measurable change in mitochondrial respiration in a non-phosphorylating state. On the other hand, we observed that mitochondria treated with a high amount of ATP (250 mM) in the presence of low GTP concentrations (5 mM) produced high levels of H_2_O_2_. Increasing GTP to 100 mM (an amount eight times the IC_50_ found in Fig. 2) leads to a marked decrease in mitochondrial H_2_O_2_ production. Thus, GTP binds mitoSUR and activates mitochondrial ATP-sensitive K^+^ entry, lowering mitochondrial ROS production withour overt changes in oxygen consumption rates.

**Figure 6.**
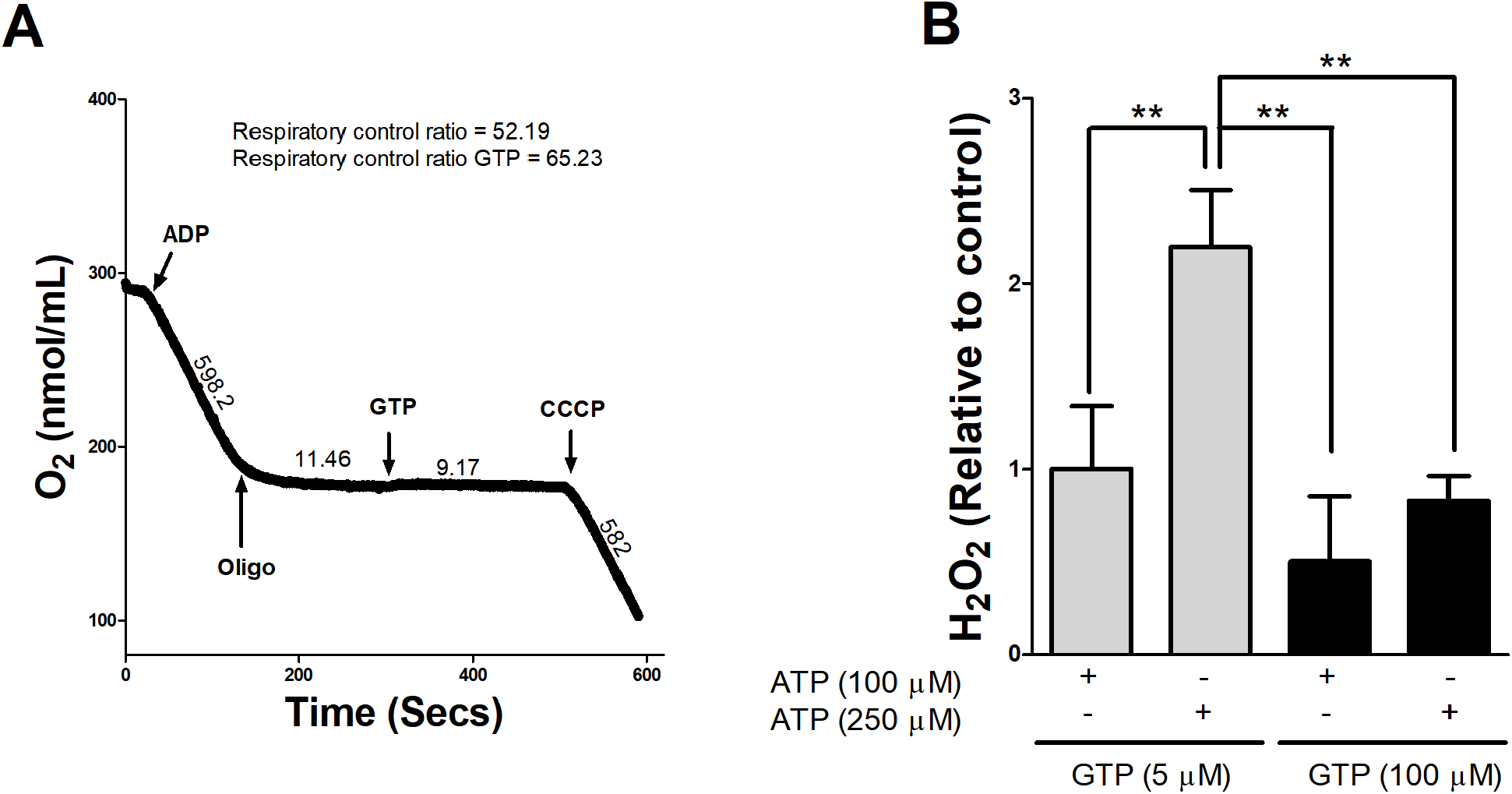
GTP-induced mitoKATP activation blocks mitochondrial ROS production. Panel A, a typical oxygen consumption trace, is shown. Mitochondria (∼0.25 mg protein/mL) were incubated at room temperature in 100 mM KCl, 10 mM HEPES, 2 mM succinate, *1 μM* rotenone, 2 mM MgCl2, 2 mM KH2PO4, pH 7.2 (KOH). Then, 1 mM ADP was added to induce state 3 respiration. 1 μg/mL oligomycin was used to induce state 4. An uncoupled state was found after the addition of 100 nM CCCP. In panel B, mitochondrial H2O2 release was measured using 50 μM Amplex Red and 1 U/mL horseradish peroxidase, in the presence of ATP (100 or 250 *μM)* and/or 100 μM GTP (5 *or 100 μM)*, as indicated. The graph depicts data from three experiments for each sample, presented as averages±S.E.M.

## 4. Discussion

This paper reports how mitoKATP senses the nucleotides ATP and GTP. The key findings are as follows: (i) Swelling assays following mitochondrial K^+^ entrance confirm that MitoKATP is blocked by ATP and opened by GTP; (ii) molecular docking analysis revealed how ATP and GTP interact with the mitoSUR protein; and (iii) pharmacological studies suggest that ATP and GTP compete for mitoSUR binding. GTP favors a mitoKATP open-state, which physiologically blocks mitochondrial ROS formation. Broadly, our work identifies how mitoKATP is regulated competitively by nucleotides (GTP and ATP). Since mitoKATP opening is a powerful way to block ischemia and reperfusion insults [7,10,17,18], a precise understanding of how nucleotides interact with mitoKATP may represent a potential target to prevent myocardial damage.

MitoKATP primarily works by transporting K+ into the mitochondrial matrix [2,15]. MitoKATP-induced mitochondrial K+ influx has attracted the interest of researchers because several lines of evidence point to this channel’s cytoprotective activity against cardiac ischemia-reperfusion [7,10,17,18] and cardiac hypertrophy [11,13]. A precise knowledge of how mitoKATP is physiologically regulated would help develop new protective strategies and explain how mitoKATP opening modulates myocardial damage. To our knowledge, this is the first study suggesting how GTP and ATP bind to mitoKATP. Actually, structural aspects and the molecular identity of this channel were very controversial, with some considering that the protein responsible for K^+^ transport or the sulfonylurea receptor were non-existent in the mitochondrial inner membrane [22]. Recently, the gene encoding a protein responsible for ATP-sensitive K+ entrance into mitochondria has been described. The same group pointed out that the mitoSUR is an ATP-binding cassette subfamily B member 8 (ABCB8) protein [2]. As a typical ABC transporter, this protein is targeted to mitochondria [23], and is composed of a transmembrane domain and a nucleotide-binding domain. As demonstrated in the crystal structure (PDB number 5OCH), this protein is a dimer that binds nucleotides (ex: ATP and ADP), and the nucleotide-binding domain faces the mitochondrial matrix [24].

As mitoKATP activity is involved in the regulation of mitochondrial volume [1], and to put our molecular docking simulation in an experimental context, we evaluated the sensitivity of mitoKATP in cardiac mitochondria to ATP inhibitory effects and GTP stimulatory effects using a swelling technique. Both techniques (docking and swelling) support the notion that nucleotide binding regulates mitoKATP leading to volume (Figs 1, and 2) and redox effects (Fig 6). Moreover, we can affirm that the ABCB8 protein (mitoSUR) binds nucleotides on the matrix side. This is also supported by the crystal structure. Indeed, early [15] and most recent [2] patch-clamp experiments demonstrated that ATP inhibits K^+^ currents when applied to the matrix side. Here, besides verifying that ATP inhibits the channel, we show that GTP opens it and propose that both nucleotides compete for the same binding site described for ADP in the ABCB8/mitoSUR protein (see crystal structure PDB number 5OCH used in this study or PDB: 7EHL). This also sheds light on the physiological regulation of the channel. It is possible that the ATP gets hydrolyzed since the nucleotide-binding domain of ATP-binding cassette proteins can bind and hydrolyze ATP [25]. In this case, we hypothesize that ADP would persist bound at the site, serving as the inhibitor molecule. Indeed, ADP inhibits mitochondrial K+ currents, although with a lower affinity [26]. Interestingly, mitoSUR/ABCB8 binds the non-hydrolyzable ATP analog (ATP-PNP) [24] but this molecule is not capable of inhibiting the mitoKATP [27]. Whether mitoSUR (ABCB8) hydrolyzes GTP or not is not clear yet.

To expand our knowledge of mitoKATP modulation by nucleotides (ATP and GTP) at the molecular level, we used molecular docking to simulate the binding of these nucleotides to ABCB8 (mitoSUR). One important functional aspect of mitoKATP activity regulation by ATP is its Mg2+ dependency, which is indispensable for the ATP inhibitory effect [28–30]. The levels of Mg^2+^ are ten times more concentrated in the mitochondrial matrix, where it complexes with both ATP and ADP [31]. Interestingly, the Mg^2+^ ion is part of the structure of ABCB8 (mitoSUR). Docking reveals that this ion interacts with the beta and gamma oxygen of the phosphate group of each nucleotide (Figs 3, 4, and 5). Our experimental results confirm the negative actions of ATP on mitoKATP opening (IC_50_ = 21.24 ± 1.4 mM) and the positive action of GTP (EC_50_ = 13.19 ± 1.33 mM), both in the presence of Mg^2+^. This study went further by suggesting a possible mechanism of action for these two nucleotides. Our docking and experimental data suggest that GTP displaces ATP-binding from mitoSUR, which suggests a competitive antagonism. So, the channel is induced to an open state since mitoSUR is a regulatory protein that binds both nucleotides. GTP action requires ATP and Mg^2+^. A mitoKATP opener (GTP) is properly tested only if the channel has previously been closed in the presence of an inhibitor. Importantly, as seen in Figure 5, increasing concentrations of ATP (100, 200, and 500 mM) lead to a shift to the right of the dose-response curves of GTP. Paucek et al. [4] tested liposome-reconstituted mitochondrial KATP channels with varying GTP concentrations in the presence of an ATP dose-response. Fig. 2 in this paper shows a shift to the right as GTP is increased from 0 to 2, 4, 8, and up to 20 μM. Our findings thus support and expand the idea promoted by Paucek and co-authors [4]. Additionally, we help illuminate the mechanism of nucleotide modulation of mitoKATP and provide insight into how and where nucleotides (ATP and GTP) potentially bind.

All the studies described and discussed above left us with a difficult question: what are the functional consequences for mitoKATP activitu of the balance between ATP and GTP under physiological conditions? MitoKATP opening leads to morphological changes (swelling, as seen in Figs 2 and 3) which play a key role in triggering pro-survival pathways, but it is difficult to reconcile how the endogenous modulation by nucleotides influences this and mitochondrial ROS production. Knowing that, we recognize that this paper may have some limitations: 1. It is difficult to up- or down-regulate nucleotide levels *in vivo* or *ex vivo*, which limits our studies since we cannot manipulate their levels, for example, in experiments involving cardiac perfusion in the Langendorff apparatus. 2. A second limitation would be the high level of ATP found in cardiomyocytes. Indeed, the ATP levels in human cardiac tissue are approximately 5.5 mM per kilogram of tissue [32]. This observation would challenge the real influence of GTP on mitoKATP activity. However, one must consider the dynamic cellular levels of ADP, GDP, and acyl CoA esterification, which also influence the channel`s activity [4]. Therefore, future studies probing for mitoKATP dynamics in the presence of the above mentioned nucleotides and experiments using modern crystallographic or cryo-EM techniques may shed light on the mechanisms of physiological or pathological mitoKATP opening and closure. To our knowledge, this is the first report demonstrating the interaction of ATP and GTP with mitoSUR.

## Conclusion

We have demonstrated the binding mechanism of nucleotides (GTP and ATP) in mitoSUR (ABCB8) of mitoKATP. GTP is an endogenous mitoKATP opener that competes with and reverses ATP inhibitory activity. The balance between GTP and ATP actions may be of further relevance via its role in modulating ROS production. These results may help in the development of new and selective mitoKATP agonists and antagonists, contributing to a better understanding of their mechanisms of action and identifying novel therapeutic targets for ischemic injury.

## Acknowledgements

The authors acknowledge Alicia Juliana Kowaltowski for critically reading the paper and for her invaluable support. Authors used UCSF chimera software for molecular graphics and analyses. UCSF Chimera was developed by the Resource for Biocomputing, Visualization, and Informatics at the University of California, San Francisco, with support from NIH P41-GM103311.

## Credit authorship contribution statement

Heberty T Facundo: Conceptualization, Investigation, Writing - original draft - review & editing, Supervision. Plinio Bezerra Palacio: Investigation, Methodology, review & editing. Geovanna Carvalho de Freitas Soares, Gabriella Moreira Bezerra Lima, and Pedro Lourenzo Oliveira Cunha: Investigation, methodology. Anna Lídia Nunes Varela: Methodology. All authors read and approved the final manuscript.

## Compliance with ethical standards

All animal experiments were approved by the Universidade Federal do Cariri animal ethics committee (protocol number 02/2020).

