## Supplementary Table 1 for "Competitive interaction between ATP and GTP regulates mitochondrial ATP-sensitive potassium channels"

**Supplementary table I:** Comparison between docked GTP and crystallized ADP showing the interaction of each residue and the residues in common (+) or not (-). The information from crystallized ADP was gathered from PDB website (code number 5OCH).

| **Residue** | **Docked GTP** | **Crystallized ADP** | **Residues in Common** |
| --- | --- | --- | --- |
| ASP244 | Van der Waals | - | - |
| ARG342 | Van der Waals | Van der Waals | + |
| TYR 481 | Van der Waals | Pi-Pi Stacked | + |
| CYS 483/  YCM 483 | Pi-Alkyl | Hydrogen Bond | + |
| ARG 484 | Pi-Alkyl | - | - |
| VAL 489 | Van der Waals | Van der Waals | + |
| GLN 508 | - | Van der Waals | - |
| SER 509 | - | Van der Waals | - |
| GLY 510 | Hydrogen Bond | Hydrogen Bond | + |
| GLY 511 | Hydrogen Bond | Hydrogen Bond | + |
| GLY 512 | Hydrogen Bond | Hydrogen Bond | + |
| LYS 513 | Attractive Charges | Attractive Charges | + |
| THR 514 | Hydrogen Bond | Hydrogen Bond | + |
| THR 515 | Hydrogen Bond | Hydrogen Bond | + |
| GLU607 | Van der Waals | - |  |
| THR 610 | Van der Waals | - | - |
| THR 611 | Van der Waals | Van der Waals | + |
