## Supplementary Fig. 1 for "Competitive interaction between ATP and GTP regulates mitochondrial ATP-sensitive potassium channels"

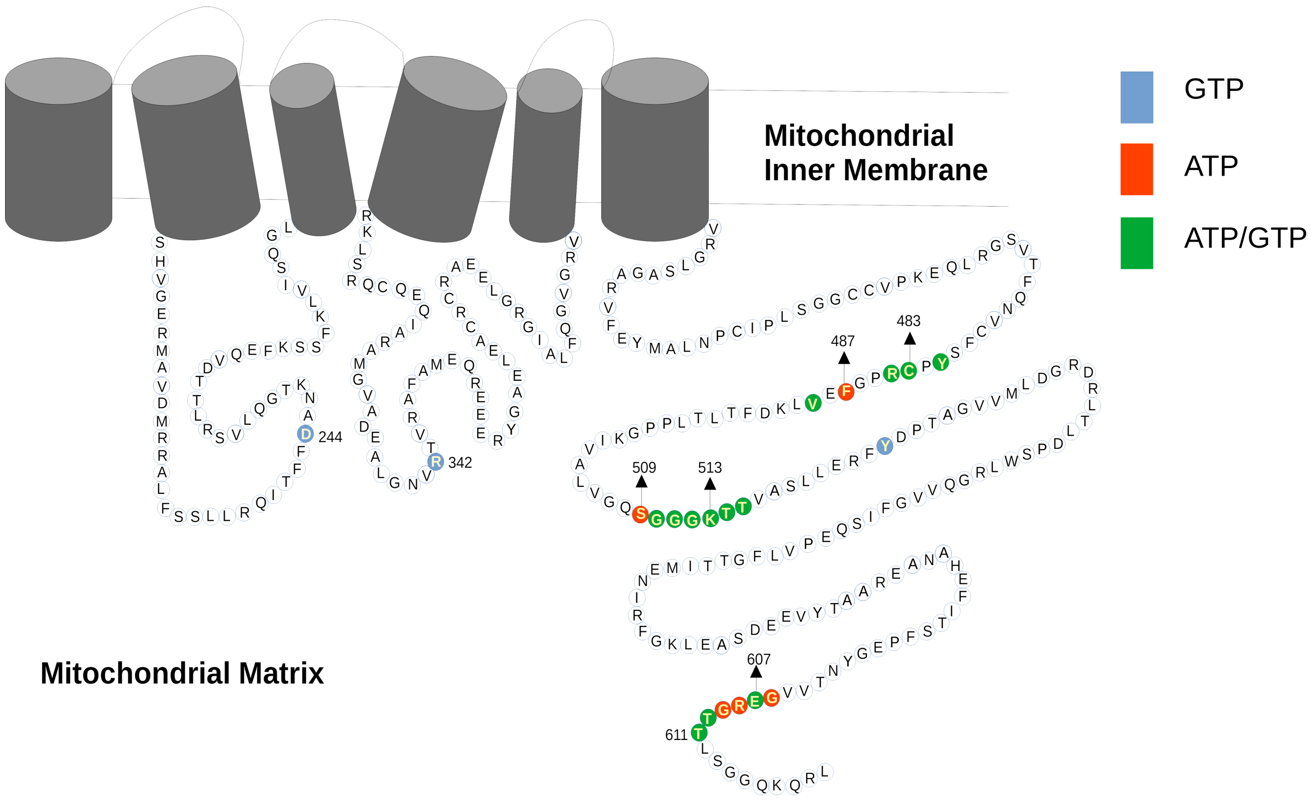


**Supplementary figure 1:** topological model shows the residues that exclusively bind GTP (Blue) and residues that exclusively bind ATP (Red). The overlapping binding residues indicating a competitive interaction for ABCB8 are colored in green.
